# A translocation-competent pore is required for *Shigella flexneri* to escape from the double membrane vacuole during intercellular spread

**DOI:** 10.1101/2024.11.11.623084

**Authors:** Julie E. Raab, Tucker B. Harju, Jody D. Toperzer, Jeffrey K. Duncan-Lowey, Marcia B. Goldberg, Brian C. Russo

**Author notes:** Address correspondence to: Brian C. Russo,.

## Abstract

Type 3 secretion systems (T3SSs) enable bacterial virulence by translocating virulence proteins (effectors) into host cells. *Shigella flexneri* require T3SS to invade and to spread between cells in the colon. In order to spread, *S. flexneri* forms membrane protrusions that push into the adjacent host cell. These protrusions are resolved into double membrane vacuoles (DMVs) that the bacteria quickly escape. The mechanisms required for escape from the DMV are poorly understood, but the T3SS translocon pore protein IpaC is essential. Here, we show IpaC forms a pore that is competent for translocation of T3SS effectors as bacteria spread between cells. To do so, we used a genetic approach to test mutations of IpaC that disrupt its ability to translocate and to form pores. We show that during spread, IpaC is efficiently inserted into the plasma membrane, the membrane-embedded IpaC forms pore complexes, and the IpaC-dependent pores translocate effectors that are necessary for *S. flexneri* to escape the DMV. We further show that T3SS activation is regulated through a distinct mechanism at spread compared to at invasion; activation of T3SS secretion does not require pore formation during spread. Thus, we show that a distinct regulation of the T3SS during *S. flexneri* intercellular spread enables the placement of effectors both around *S. flexneri* and across membranes of the DMV. Altogether, this study provides new insights into how *S. flexneri* escapes the DMV.

**IMPORTANCE:** The type 3 secretion system (T3SS) is required for virulence in many bacterial pathogens that infect humans. The T3SS forms a pore through which virulence proteins are delivered into host cells to enable bacterial infection. Our work investigates the *Shigella* translocon pore protein IpaC, which is essential not only for bacteria to invade cells, but also for bacteria to spread between cells. An ability to spread between cells is essential for pathogenesis, thus understanding the mechanisms that enable spread is important for understanding how *S. flexneri* infection causes illness. We show that IpaC delivers virulence factors across the host membrane for *S. flexneri* to efficiently spread. This study furthers our understanding of the mechanisms involved in T3SS secretion and of translocon pore function during *S. flexneri* intercellular spread.

## INTRODUCTION

*Shigella flexneri* is a gram-negative bacterial pathogen and a major cause of moderate-to-severe diarrheal illness in humans^1^. Disease arises from invasion of and spread between epithelial cells in the colonic epithelium. Intercellular spread causes death of more epithelial cells than invasion alone, and this disruption of the intestinal barrier increases colonic inflammation^2–4^. Thus, the ability to spread is essential for disease progression to severe, bloody diarrhea^5, 6^. Both invasion and spread require a type 3 secretion system (T3SS). The T3SS is a needle-like structure that delivers virulence proteins (effectors) into epithelial cells. The delivered effectors reprogram cellular pathways that are necessary for *S. flexneri* to establish an intracellular niche.

To access the host cytosol, the effectors of *S. flexneri* are delivered across the plasma membrane through a pore, known as the translocon pore. This pore is generated by T3SS-mediated delivery of the bacterial proteins IpaC and IpaB into the plasma membrane^7^. The translocon pore protein IpaC then interacts with host cytosolic intermediate filaments to activate secretion of effectors through the T3SS into the host cell^8, 9^. Thus, at invasion, IpaC forms a pore that regulates the secretion of effector proteins through the T3SS. In contrast to invasion, intermediate filaments are dispensable to activate T3SS secretion when *S. flexneri* spreads^10^. IpaC enables cytosolic *S. flexneri* to form protrusions in the host membrane by releasing tension at the membrane through its interaction with β-catenin^10, 11^, a cytosolic protein and component of adherens junctions^12, 13^. The protrusion extends into and is engulfed by the neighboring cell^14^. The engulfed protrusion resolves into a double membrane vacuole (DMV), which the bacteria escape into the cytosol of the neighboring cell. Whereas mutations of IpaC that specifically disrupt its interaction with β-catenin reduce the efficiency of spread by limiting protrusion formation^10^, the complete loss of IpaC prevents spread^7, 15^. Bacteria lacking IpaC are observed as trapped in the DMV^15, 16^, suggesting that IpaC has additional functional activity during spread that is essential for escape from the DMV; this functional activity of IpaC during escape from DMV has not been characterized.

The objective of this study was to determine how IpaC contributes to escape from the DMV. We show that, in the absence of IpaC, secretion of type 3 effectors was efficient, but more bacteria were retained in DMVs during intercellular spread. To assess how IpaC contributes to escape from the DMV and intercellular spread, we developed a system that enabled the study IpaC variants with specific functional defects. Using this system, we observed that the formation of T3SS translocation-competent pores was required for bacteria to spread. Pore formation alone was not sufficient; it is necessary that pores are competent for translocation of effectors to facilitate escape by *S. flexneri* from the DMV and spread between cells. Overall, these data show that *S. flexneri* must produce a pore that is competent for translocation so as to enable bacterial escape from DMVs and promote intercellular spread.

## RESULTS

### IpaC and IpaB are present in the host membrane during spread

IpaC associates with the cytosolic adherens junction protein β-catenin^10, 11^, but whether IpaC is positioned in the membrane at spread is uncertain. The function and location of IpaC are well characterized during invasion; IpaC is in the host membrane as part of the translocon pore, and its C-terminal region is within the host cytoplasm^7^. To determine whether translocon pore proteins associate with the membrane during spread, we tested whether IpaC and IpaB were localized to the host membrane throughout the course of infection. We infected HeLa cells with *S. flexneri* Δ*ipaC* expressing *ipaC* under the control of the arabinose-inducible pBAD promoter (*Sf* Δ*ipaC*::*ipaC*). To examine the abundance of our target proteins in the host membrane, infected cells were fractionated at the indicated times using the detergent saponin to permeabilize the plasma membrane of the infected cells and to recover the cytosolic fraction. Solubilization of the membranes by the detergent triton X-100 enabled recovery of the membrane components. The insoluble components and intact bacteria were removed by centrifugation, as done previously^8–10, 17–20^. We show that, for spread, IpaC presence in the membrane requires its continuous production. In the absence of arabinose, the abundance of IpaC is significantly reduced in the host membrane fraction and from the bacteria by 2.5 hours of infection (Fig. 1A-B). Similar to invasion, the abundance of IpaB in the membrane was independent of the abundance of IpaC (Fig. 1A and C), which shows the insertion of IpaB into the membrane at spread is also independent of IpaC presence. To exclude the possibility that new invasion events were the source of the IpaC and IpaB in the membrane during spread, gentamicin was added after 45 minutes of infection to kill remaining extracellular bacteria. In these experiments, the presence of the bacterial cytosolic protein GroEL only in the bacterial fraction indicated *S. flexneri* was not lysed by our fractionation approach. The eukaryotic plasma membrane protein caveolin-1 showed the plasma membrane components were in the expected fraction. Similar to our observations in HeLa cells, translocon pore proteins localized to the membrane of Caco-2 cells, a polarized human colonic epithelial cell line (Fig. S1). Altogether, these data show the translocon pore proteins IpaC and IpaB are continuously inserted into the plasma membrane during spread, regardless of cell type and polarity.

**Figure 1:**
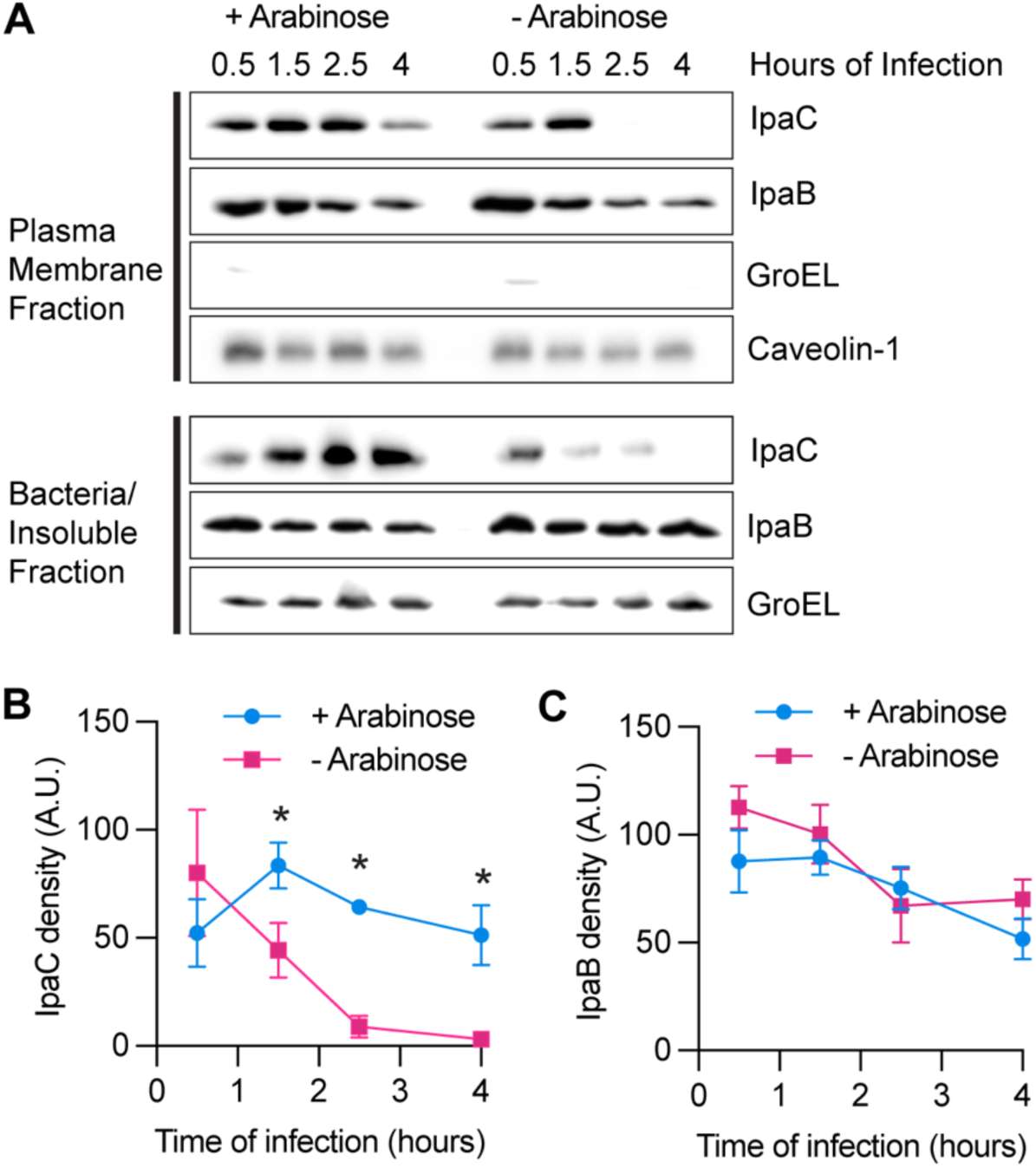
IpaC and IpaB are present in the host membrane during spread. **A)** Representative western blots showing abundance of IpaC, IpaB, GroEL (bacterial cytoplasmic protein), and Caveolin-1 (eukaryotic membrane protein) in the membrane fraction (top panels) and abundance of IpaC, IpaB, and GroEL in the insoluble fraction/bacteria (bottom panels). Quantitative measurements of IpaC (**B)** or IpaB (**C)** in the membrane fraction with (light blue) or without (magenta) arabinose included in the media to induce IpaC production. **B & C)** At least two independent experiments were performed for each infection condition at each timepoint. Data are mean ± SEM, some error bars are smaller than the symbol and are not visible. Mixed effects analysis with Fisher’s multiple comparisons test, *p<0.05.

### IpaC contributes to bacterial escape from DMVs, but secretion is activated independent of IpaC

During invasion, T3SS secretion is triggered by the interaction of IpaC with intermediate filaments^8, 9^. Intermediate filaments are dispensable during spread, and the secretion of T3SS effectors is activated in their absence and is associated with the interaction of bacteria with the plasma membrane^10, 16^. Since IpaC interaction with intermediate filaments activates secretion during invasion, we asked whether T3SS activation still requires an IpaC-dependent mechanism during spread. We infected HeLa cells with bacteria that produced or lacked IpaC and examined T3SS activation during spread. HeLa cells produced a membrane-anchored YFP^21^, which enabled visualization of bacteria in a protrusion or DMV. *Sf* Δ*ipaC*::*ipaC* regulate IpaC production from the pBAD promoter and IpaC abundance can by regulated by the presence of arabinose. These bacteria also encoded a fluorescent reporter of T3SS secretion (pTSAR), which constitutively produces mCherry and inducibly produces GFP when the T3SS effector OspD1 is secreted^16^. Here, we used secretion of OspD1 as a general indicator of T3SS activation^8–10, 16, 19^. Spread starts around 2 hours of infection, during which bacteria form protrusions that resolve into double membrane vacuoles (DMVs)^22^; we examined infection at 4 hours when protrusion formation is robust and IpaC is lost (Fig. 1). As anticipated from previous studies^15, 16^, the percentage of bacteria present in DMVs increased when the bacteria did not produce IpaC (Fig. 2B and Table S1), which confirmed IpaC is required for efficient escape from DMVs. During invasion, the activation of T3SS secretion requires both pore formation in the plasma membrane and an activation step that is dependent on IpaC interaction with intermediate filaments. In contrast to invasion, T3SS activation was efficient at 4 hours of infection even for bacteria that lacked IpaC and were unable to form pores (Fig. 2C and Table S1), which showed effector secretion can occur in the absence of IpaC and pore formation.

**Figure 2:**
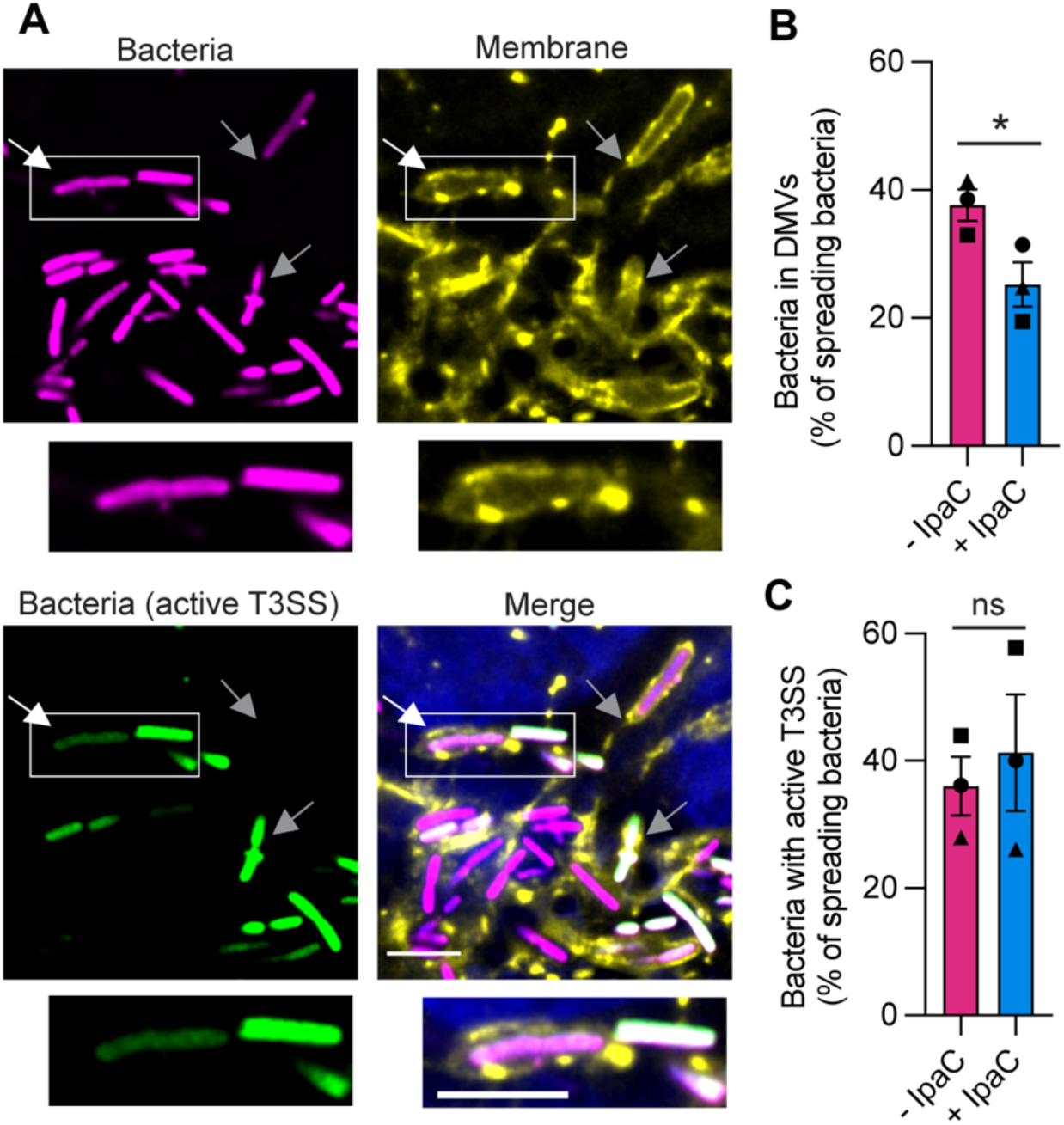
IpaC contributes to bacterial escape from DMVs, but secretion is activated independent of IpaC. **A)** Representative immunofluorescence images of HeLa pmbYFP cells infected with bacteria for 4 hours. Bacteria in a DMV (white arrow) or protrusion (grey arrow). White rectangle: zoomed in image of bacterium within a DMV (below). Magenta, all bacteria; green, bacteria with active T3SS; yellow, HeLa plasma membranes; blue, DNA; scale bar, 5 μm. **B & C)** Percentage of bacteria that are spreading (present in DMV or protrusion) that are in DMVs (**B)** or that have active T3SS (**C)** from images represented in **A**. Bacteria producing (+IpaC, aqua) or not producing (-IpaC, magenta) IpaC during infection. Data are mean ± SEM of three independent experiments, each experiment matched by symbol. ns (not significant), *p<0.05 by paired t-test.

Since T3SS activation is associated with membrane contact during spread^16^ and more bacteria lacking IpaC remain in close contact with the plasma membrane of the DMV, we also observed more bacteria produced GFP in the absence of IpaC (Fig. S2 and Table S1). This indicated that the bacteria trapped in the DMV maintain active T3SS for a longer duration than the bacteria with IpaC that quickly escape the DMV. These data show the T3SS is activated in a distinct manner at invasion compared to at spread, and they support previous observations that repeated membrane contact is associated with activation of T3SS^16^. Altogether, these data show that IpaC is required for *S. flexneri* to escape the DMV and that IpaC is dispensable for T3SS secretion during spread.

### The formation of translocation-competent pores is essential for spread

Despite not regulating T3SS effector secretion, we sought to test whether other known activities of IpaC were required during intercellular spread. We speculated that the ability of IpaC to form a pore would still be required to lyse the membranes and to escape from the DMV. Similarly, pore formation enables escape from the vacuole associated with bacterial invasion^23^. To test this, we made use of mutations that disrupt the ability of IpaC to form a functional pore. Loss of the C-terminus disrupts the interactions that enable docking of the T3SS needle to the T3SS pore, the ability to interact with β-catenin, and the translocation of effectors^9–11, 24^. Deletion of and mutations around the coiled-coil region of IpaC disrupt the ability of the pore to undergo conformation changes that enable translocation of effectors from *S. flexneri* into the host cytosol^19^. Because some mutations prevent T3SS-mediated effector translocation and invasion, we created *S. flexneri* strains that produce an inducible version of IpaC, which enables invasion and whose abundance can become undetectable by spread. A second version of IpaC is also produced that is constitutively expressed to study the effect of IpaC mutations on intercellular spread. By including a 1X FLAG tag on the inducible version of IpaC at amino acid position 37 (IpaC- FLAG), we can differentiate it from the constitutive IpaC by a size shift on western blot (Fig. S3A). IpaC-FLAG is similar in functionality to wild-type IpaC as it can complement *S. flexneri ΔipaC* to form pores and enable effector translocation (Fig. S4). Plasmids encoding the inducible IpaC-FLAG and constitutive IpaC variants were transformed into *S. flexneri ΔipaC,* and resulting strains were tested for IpaC production and secretion. T3SS secretion remained regulated by Congo red in the presence of IpaC variants and IpaC-FLAG (Fig. S3), which demonstrated that these mutations neither affect the ability of IpaC to be secreted nor the activation of the T3SS in general.

Previous investigation of these IpaC variants shows they insert in plasma membranes^8, 9, 19^, but IpaC Q308P forms smaller pores that lyse membranes less efficiently and IpaC Δcoiled-coil is unable to lyse membranes compared to IpaC WT^19^. Additionally, only WT IpaC and IpaC A354P translocate effectors during invasion, with IpaC A354P showing reduced translocation compared to IpaC WT^19^. We characterized the ability of these IpaC variants to enable intercellular spread using a plaque assay. In this assay, the number of plaques correlates with the efficiency of invasion and the area of plaques is associated with the efficiency of intercellular spread. We induced production of IpaC-FLAG prior to invasion so that strains could invade cells regardless of which constitutive version of IpaC was produced. Plaques were not detected from strains lacking IpaC nor from strains producing IpaC Q308P, IpaC ΔC-terminus, or IpaC Δcoiled-coil. In contrast, large plaques were formed by strains producing WT IpaC or IpaC A354P (Fig. 3A-B). Since plaques were observed only for strains producing versions of IpaC that could both form pores and translocate effectors, these findings indicate that spread requires pores that are sufficiently open such that effectors can transit through them. In contrast, when IpaC-FLAG production is maintained throughout the duration of the assay, all strains can form plaques, which showed that the defects in plaque formation observed for particular IpaC variants were not the result of the strain being unable to invade cells initially, but rather, due to the inability of specific constitutively produced IpaC mutants to support intercellular spread (Fig. 3C-D). Overall, these data suggest that IpaC is required for the formation of a translocation-competent pore to enable bacteria to efficiently spread.

**Figure 3:**
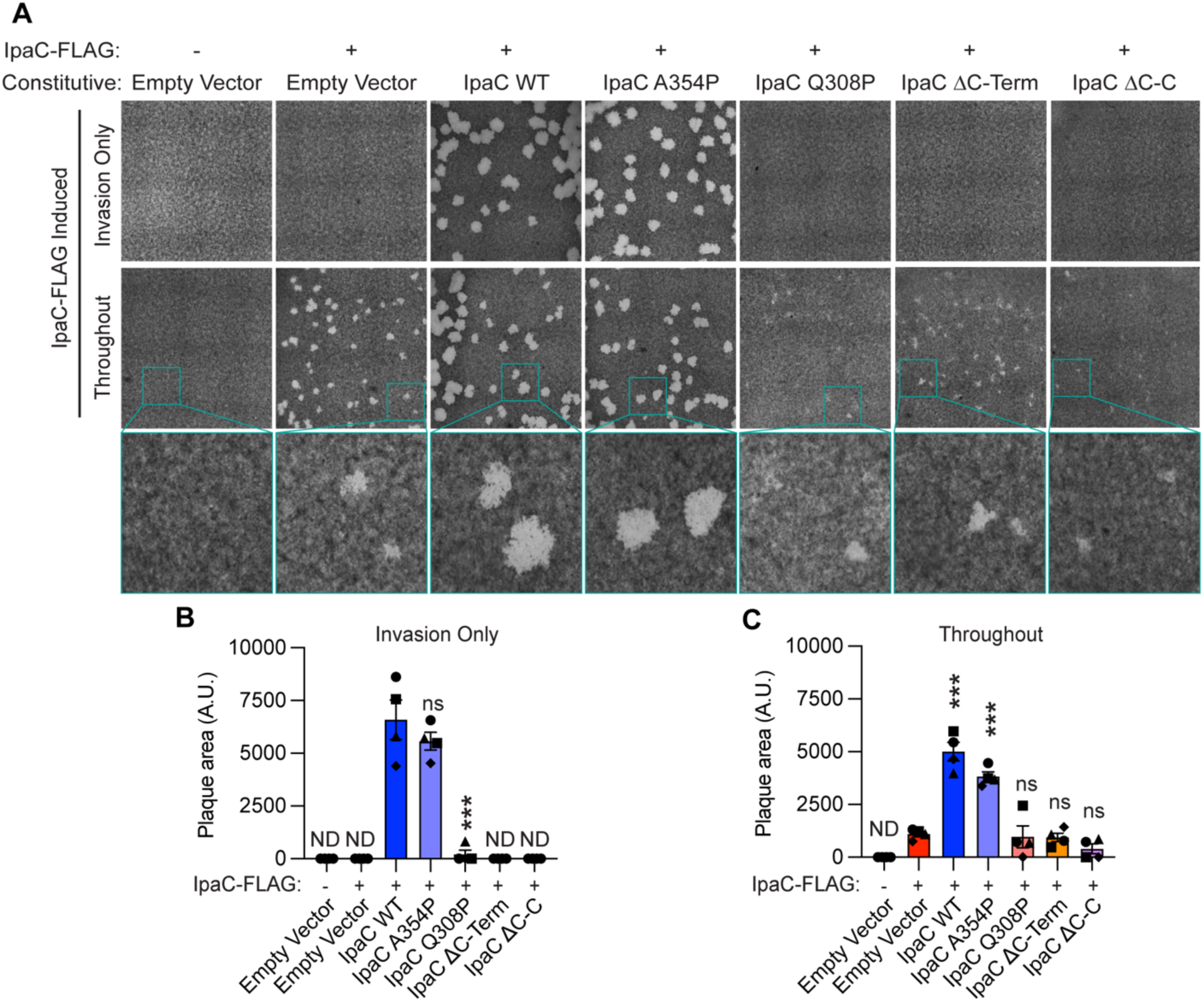
The formation of translocation-competent pores is essential for spread. **A)** Representative images of plaque assays after 48 hours of infection of MEF cells. Top: IpaC-FLAG induced only before infection to enable invasion. Middle: IpaC-FLAG induced throughout the duration of infection. Bottom: Zoomed in from teal square in middle panels. C-Term (C-terminus), C-C (Coiled-coil). **B & C)** Quantification of plaque areas from infections where IpaC-FLAG was induced **(B)** for invasion only or **(C)** throughout the infection from images represented in A. Data are mean ± SEM of four independent experiments, each experiment matched by symbol. One-way ANOVA with Dunnett’s multiple comparisons; ND (no detected plaques), ns (not significant), ***p<0.001.

### Insertion of pore proteins into the host membrane is not sufficient for spread

To further determine how the formation of translocation-competent pores is required for intercellular spread, we first identified whether all variants of IpaC were inserted into the host membrane at spread. We induced production of IpaC-FLAG only during the back dilution so that strains could invade cells regardless of which IpaC variant was produced. We then infected cells with these strains and fractionated infected cells at 4 hours of infection as described above. For all strains tested, constitutively produced IpaC variants were inserted into the plasma membrane at 4 hours of infection (Fig. 4A-B). As expected, in the absence of induction, IpaC-FLAG is not detectable at 4 hours of infection (Fig. 4A). IpaB is detected in the membrane at spread regardless of which IpaC variant is present in the membrane (Fig. 4A-C). Together, these data show that the inability of strains producing IpaC Q308P, IpaC ΔC-terminus, or IpaC Δcoiled-coil to spread is not due to their failure to insert these IpaC variants into the host plasma membrane during spread. Instead, the failure to spread is likely attributable to other functional defects associated with these IpaC variants such as their ability to support pore formation and translocation.

**Figure 4:**
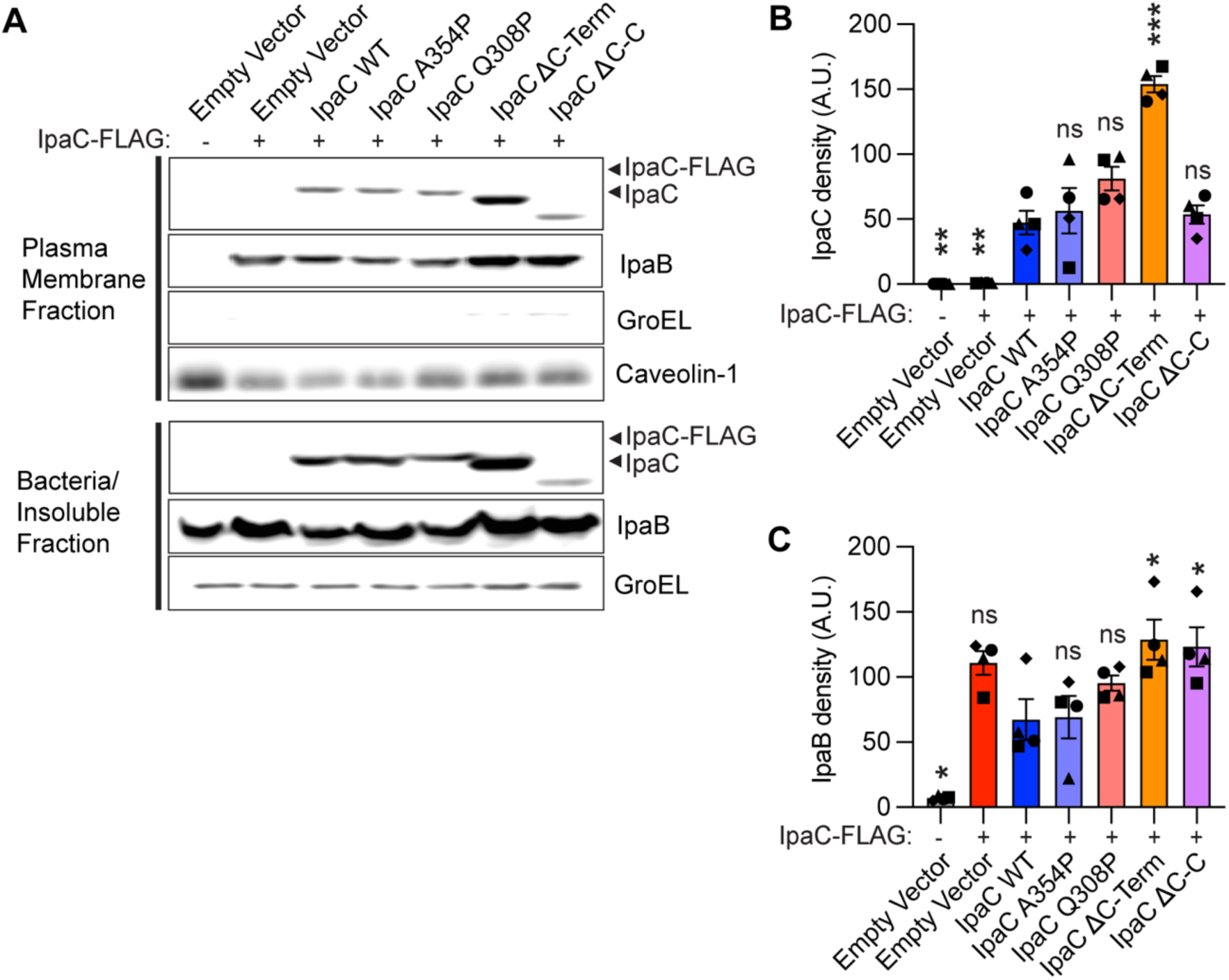
Insertion of pore proteins into the host membrane is not sufficient for spread. **A)** Representative western blots showing abundance of IpaC variants, IpaB, GroEL, and Caveolin-1 in the membrane fraction (top panels) and of IpaC, IpaB, and GroEL in the bacteria/insoluble fraction (bottom panels). IpaC-FLAG was only induced before infection to enable invasion. C-Term (C-terminus), C-C (Coiled-coil). **B & C)** Quantitative measurements of IpaC **(B)** or IpaB **(C)** in the membrane fraction. Data are mean ± SEM of four independent experiments, each experiment matched by symbol. ns (not significant), *p<0.05, **p<0.01, ***p<0.001 by one-way ANOVA with Dunnett’s multiple comparisons test.

### Translocation-competent pores are required for *S. flexneri* to recruit Rac1 and to escape from DMVs

T3SS effectors such as IcsB and IpgB1 have functions in the recipient host cytosol that contribute to escape from the DMV^21, 25–32^. IpgB1 regulates Rho family GTPases and recruits Rac1 around DMVs in a manner that enables bacterial escape from DMVs^30–33^. How effectors function across the DMV membrane is unclear. It is presumed there may be translocation, but it is unclear whether the T3SS translocon pore functions to translocate effectors for several reasons. 1. There are two plasma membranes present in the DMV, and it is unknown how the pore might mediate effector translocation across these membranes; 2. As we show above, the regulation of T3SS-mediated secretion is uncoupled from pore formation, which suggests translocation may not be possible. We sought to investigate the relationship between effector function in the donor cell and the ability to form translocation-competent translocon pores. First, we confirmed that the effectors IcsB and IpgB1 are necessary for efficient escape from DMVs and that the recruitment of Rac1 to DMVs is dependent on the expression of *ipgB1.* Similar to previous investigations^30^, we infected HT-29 intestinal epithelial cells with WT *S. flexneri, S. flexneri ΔicsB* or *S. flexneri ΔipgB1* and at 5 hours of infection, we observed more *S. flexneri ΔicsB* and *S. flexneri ΔipgB1* in DMVs than WT *S. flexneri* (Fig. S5A-B and Table S2). As determined prior^30^, we also observed Rac1 was recruited to fewer DMVs in cells infected with *S. flexneri ΔipgB1* (Fig. S5A and C, and Table S2). This was not simply due to the amount of time spent in the DMV but specific to the absence of IpgB1; *S. flexneri ΔicsB* are delayed in escape from DMVs similar to *S. flexneri ΔipgB1*, but *S. flexneri ΔicsB* efficiently recruit Rac1 to DMVs (Fig. S5 and Table S2). We additionally confirmed these observations in HeLa cells (Fig S6 and Table S3). Thus, Rac1 colocalization with DMVs requires IpgB1, and Rac1 recruitment is not diminished due to less efficient DMV escape.

We next tested whether the formation of translocation-competent pores was associated with bacterial escape from DMVs and recruitment of Rac1 to DMVs. We infected HT-29 cells with *S. flexneri ΔipaC* producing an inducible IpaC-FLAG and an empty vector or constitutive version of IpaC (WT IpaC, IpaC A354P, or IpaC Q308P). We induced production of IpaC-FLAG only during the back dilution so that all strains could invade cells. The percentage of bacteria in DMVs and the number of DMVs associated with Rac1 was quantified by florescence microscopy for each strain of bacteria (Fig. 5A). Bacteria producing IpaC Q308P, which forms a pore that is too narrow to translocate effectors, and bacteria that harbor an empty vector, which are unable to form a pore and do not translocate effectors, were more likely to be present in DMVs than bacteria producing WT IpaC or IpaC A354P, which can form translocation-competent pores (Fig. 5B and Table S4). While all strains described express *ipgB1* and thus have the capacity to secrete IpgB1 to recruit Rac1, we observed that only bacteria producing WT IpaC or IpaC A354P showed recruitment of Rac1 around DMVs (Fig. 5C and Table S4), which shows that mutations in IpaC that prevent translocation block Rac1 recruitment. Similar observations were made in HeLa cells infected with these strains (Fig S7 and Table S5). Overall, these data show that effector translocation through translocon pores is required for effector activity and escape of *S. flexneri* from DMVs.

**Figure 5:**
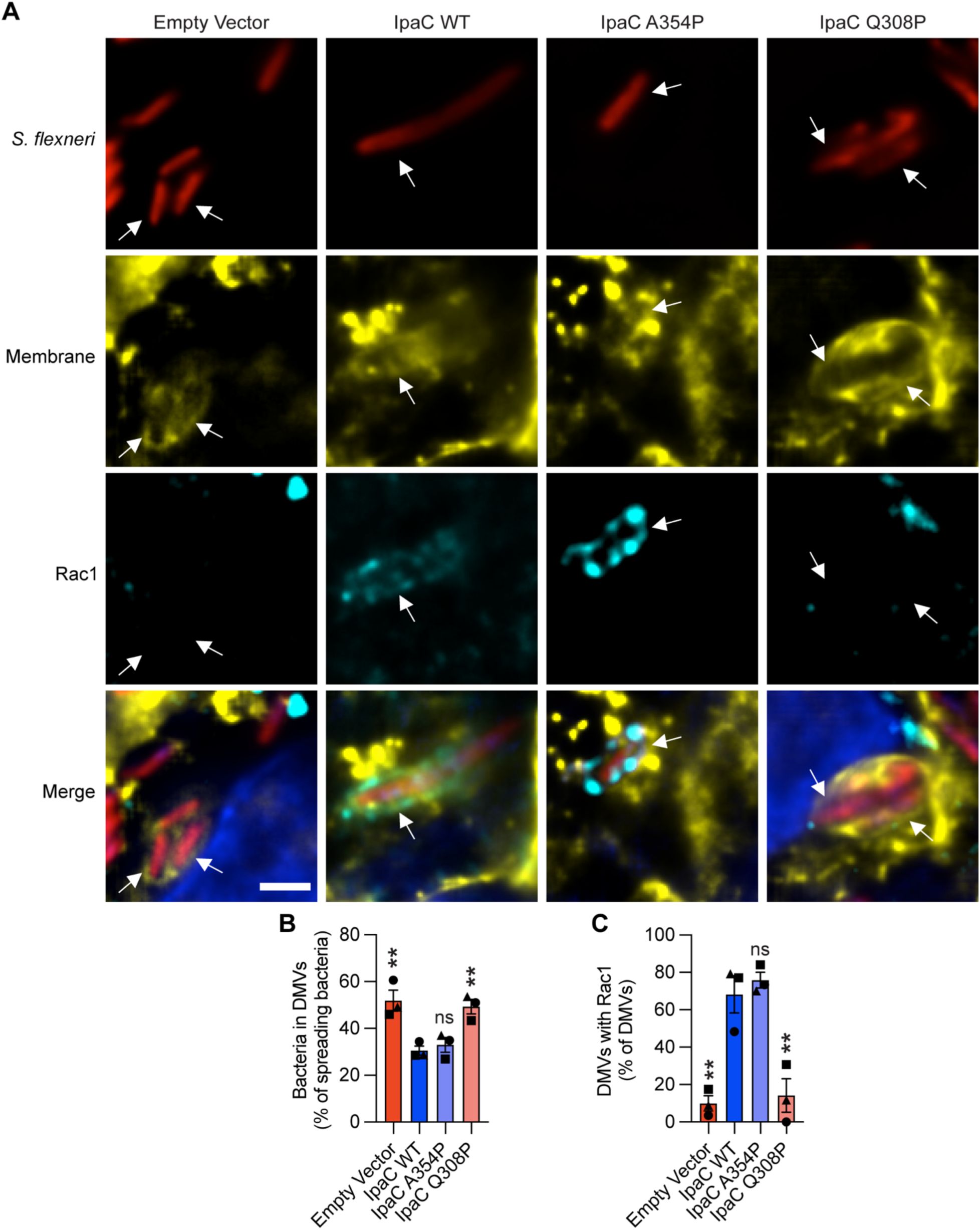
Translocation-competent pores are required for *S. flexneri* to recruit Rac1 and to escape from DMVs. **A)** Representative immunofluorescence images of HT-29 pmbYFP cells infected with bacteria producing indicated IpaC variant at 5 hours of infection. All strains included IpaC- FLAG and were induced only before infection to enable invasion. Bacteria within a DMV (white arrow). Red, bacteria; yellow, HT-29 cell membranes; cyan, Rac1; blue, DNA; scale bar, 2 μm. From images represented in **A,** percent of bacteria that are spreading and in DMVs **(B)**, and percent of bacteria within DMVs that colocalize with Rac1 **(C)**. **B-C)** Data are mean ± SEM of three independent experiments, each experiment matched by symbol. ns (not significant), **p<0.01 by one-way ANOVA with Dunnett’s multiple comparisons test.

## DISCUSSION

Intercellular spread is essential for the virulence associated with *S. flexneri* infections. IpaC is essential for intercellular spread, but the functional requirements of IpaC at this stage of infection are uncertain. Here, we built upon previous investigations of IpaC function during *S. flexneri* invasion of epithelial cells to characterize IpaC function during intercellular spread and escape from DMVs. We show new evidence that T3SS pore proteins are inserted into the host plasma membrane during spread and that a translocon-competent pore enables effector function in the neighboring cell for *S. flexneri* to efficiently escape the DMV.

Our data show that a pore capable of translocating effectors is necessary for intercellular spread. How translocation might occur in light of the regulatory differences observed in T3SS activity at spread and at invasion is unclear. Typically, T3SS-mediated translocation occurs when the effector proteins are loaded into the T3SS base in the bacterial cytosol and then transported through the T3SS needle and across the T3SS translocon pore in a single step. This is facilitated by a close interaction, known as docking, of the needle and the pore resulting from the interactions of the translocon pore protein IpaC with intermediate filaments in the host cell^8^. Since intermediate filaments are dispensable for *S. flexneri* during spread^9^, bacteria may not need to dock, and so translocation may not occur as a single step at spread. In support of this, a mutant that is unable to support docking, IpaC R326W, efficiently activates secretion^10^, and we show that the presence of IpaC is not required to activate secretion at spread (Fig. 2C). There is evidence that *Yersinia* T3SS effector translocation can occur in the absence of docking by using a two-step process in which effectors are first secreted through the T3SS needle into the space outside the bacterium, and in a second step, effectors are transported through the T3SS translocon pore into the host cytosol^34^. We anticipate that a two-step translocation process would be beneficial to bacteria that are spreading. Effectors necessary for protrusion formation would already be secreted outside of the bacterium and thus appropriately localized in the cytosol of the donor cell, where they function to resolve protrusions. In contrast, bacteria only able to utilize one-step translocation would likely only translocate effectors into the recipient cell or into the extracellular space between host cells and would not be able to function to initiate or resolve protrusions from within the donor cell. Equally possible is that the contact with the donor membrane generates the formation of a pore that is unable to interact with the T3SS needle and leads to activation of secretion with or without the ability of effectors to pass through the pore; this may allow both the pore and the secreted effectors to permeabilize the initial plasma membrane (Fig. 6). Subsequent contact with the recipient plasma membrane by the T3SS would be anticipated to function similar to invasion in which the bacterium is located at the outer leaflet of a plasma membrane and one-step translocation would occur (Fig. 6). In either case our model is speculative and further investigation is required to determine how the T3SS translocon pore supports effector secretion and translocation during spread.

**Figure 6:**
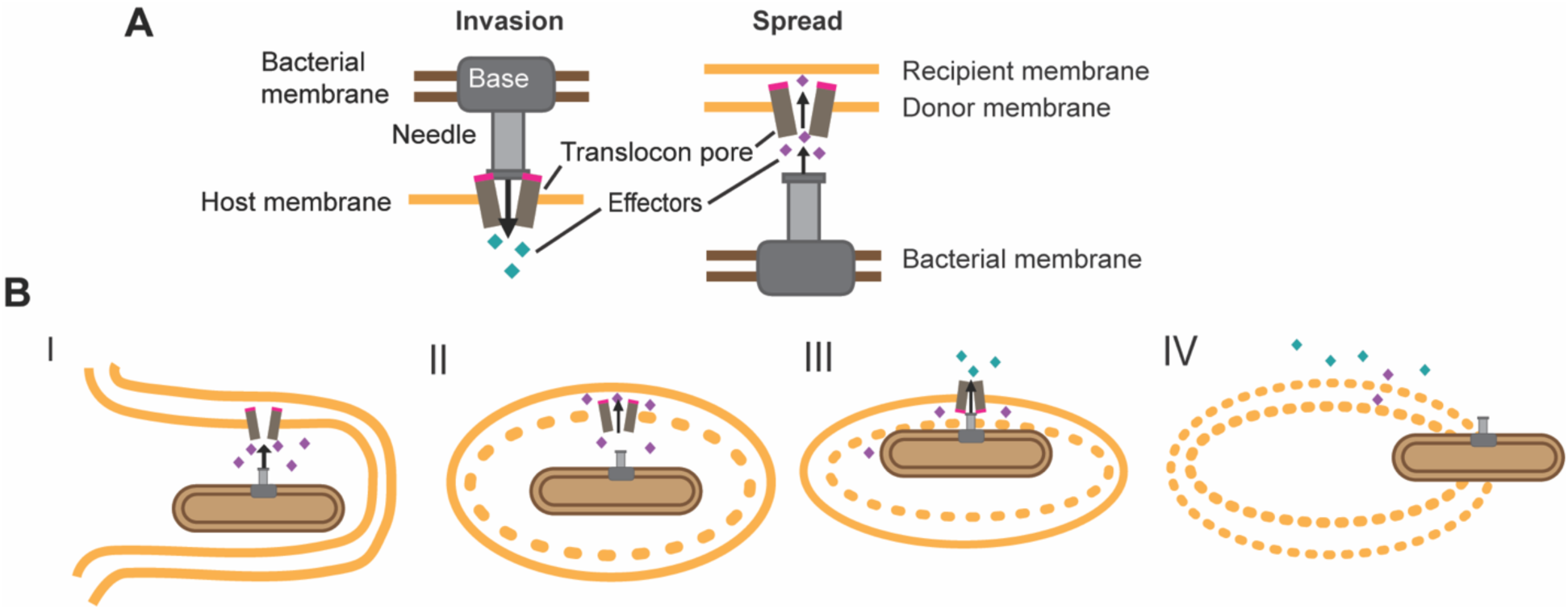
Model of translocation during *S. flexneri* infection. **A)** Schematic of effector translocation across the host membrane in a one-step process at invasion (left) and two-step process during spread (right). Teal diamonds represent effectors translocated in a one-step process and purple diamonds represent effectors translocated in a two-step process. **B)** Model of how effectors may be translocated across the DMV during spread. **I.** Upon contact with the host membrane, *S. flexneri* secretes effectors and forms a protrusion. **II.** Insertion of the pore enables effectors to translocate across the donor membrane of the DMV. Pore insertion and translocation of effectors likely permeabilize the donor membrane. **III.** Contact with the outer leaflet of the recipient membrane could enable insertion of the pore similar to invasion and result in one-step translocation. **IV.** Effectors translocated into the cytosol of the recipient cell enable bacterial escape from the DMV.

Previous studies show that insertion of IpaC and IpaB are sufficient to lyse the entry vacuole at invasion^23, 35^, suggesting that the ability to form pores at spread could enable escape from the DMV. Interestingly, IpaC Q308P can partially lyse the membranes of red blood cells^19^, showing it forms a partially open pore, but *S. flexneri* producing IpaC Q308P was unable to escape DMVs (Fig. 5B), which showed that formation of a pore is not sufficient to escape from the DMV or that a larger pore is required. Interestingly, variants of IpaC that could not form pores and were likely unable to translocate effectors showed greater defects in the ability to recruit Rac1 to the DMV than bacteria that were missing just *ipgB1*, suggesting that IpgB1 may function with additional translocated effectors to mediate Rac1 recruitment around the DMV (Fig 5C and S5C). Overall, these data show that translocon pore formation on its own is not sufficient to mediate escape from the DMV and that effector translocation is important.

During invasion, *S. flexneri* interacts initially with outer leaflet lipids of the mammalian membrane, but during spread, *S. flexneri* is in the cell cytosol, from where the T3SS initially interacts with the inner leaflet of the membrane. Both leaflets contain cholesterol, but the outer leaflet maintains a net neutral charge, whereas the inner leaflet has a net negative charge^36, 37^. Our data indicate that the regulation of effector secretion at spread is distinct from invasion. We thus speculate that the negative charge of the inner leaflet lipids may trigger secretion of effectors from *S. flexneri* and that proximity of the T3SS needle to the membrane allows the pore proteins IpaC and IpaB to insert without diffusing far from the target membrane.

In our studies, there were differences in how the IpaC variants were secreted in response to Congo red or in response to infection. We observed that IpaC ΔC-terminus was more abundant in the membrane compared to the other variants of IpaC (Fig. 4B), but it was not secreted as efficiently as WT IpaC when Congo red was used to activate T3SS secretion (Fig. S3B). We also noticed that, whereas IpaC Δcoiled-coil had significantly lower abundance in the supernatant of Congo red secretion, the abundance of IpaC Δcoiled-coil in the membrane was similar to other IpaC variants (Fig. 4B and S3B). We envision two possible reasons for this apparent difference between the amount of these IpaC variants secreted in response to Congo red and the amount detected in the membrane. 1. This difference could perhaps indicate that IpaC ΔC-terminus or IpaC Δcoiled-coil are more stable when inserted into a membrane and less prone to degradation than when present in the supernatant. 2. The deletion of the last 15 amino acids in the IpaC ΔC-terminus variant or the 35 amino acid deletion in the IpaC Δcoiled-coil variant alters the isoelectric point of the protein, perhaps favoring integration into the host membrane.

Overall, our results show that the escape of *S. flexneri* from DMVs requires translocation-competent pores. Since intercellular spread is essential for *S. flexneri* pathogenesis, it is important to understand the distinct regulation and function of the *S. flexneri* T3SS and associated pore proteins during infection. These studies provide insights into how *S. flexneri* may regulate the placement of effectors to carry out functional activities both around the bacteria and across membranes.

## MATERIALS AND METHODS

### Bacterial Strains and Culture

All *Shigella* strains used in this study are isogenic derivatives of *Shigella flexneri* strain 2457T (Table 1). *S. flexneri* strains were cultured in trypticase soy broth with appropriate antibiotics at 37°C at 250 rpm. *ipaC* or *ipaC-FLAG37* was cloned under control of the pBAD promoter on the pBAD33 plasmid. An *ipaC* variant (WT, A354P, Q308P, ΔC- terminus, or Δcoiled-coil) was cloned into a plasmid under the control of a constitutively active promoter; this plasmid also included mCherry under control of the rpsM promoter. We named this plasmid pMEIC (<u>m</u>Cherry <u>e</u>xpressing*, ipaC* <u>c</u>onstitutive). Cloning and plasmid preparation was performed within *E. coli* DH10B, and plasmids were selected with appropriate antibiotics in Luria Broth. Plasmids were transformed into *S. flexneri ΔipaC* and selected with appropriate antibiotics. Inducible production of IpaC or IpaC- FLAG was enabled by the regulation of IpaC from the *pBAD* promoter. Constitutive production of IpaC variants was enabled by the *ipaH7.8* promoter.

**Table 1:** List of strains used in this study.

| Bacterial Strain | Plasmid 1 | Plasmid 2 | Plasmid 3 | Source | Reference |
| --- | --- | --- | --- | --- | --- |
| *S. flexneri* 2457T ΔipaC | | | | Laboratory stock | [9] |
| *S. flexneri* 2457T ΔipaC | pBAD33-ipaC | | | Laboratory stock | [9] |
| *S. flexneri* 2457T ΔipaC | pBAD33-ipaC-FLAG37 | | | This study | This study |
| *S. flexneri* 2457T ΔipaC | pBAD33-ipaC | pBR322-Afa-1 | | Laboratory stock | [17] |
| *S. flexneri* 2457T ΔipaC | pTSAR | | | Laboratory stock | [9] |
| *S. flexneri* 2457T ΔipaC | pBAD33-ipaC | pTSAR | | Laboratory stock | [9] |
| *S. flexneri* 2457T ΔipaC | pBAD33-ipaC-FLAG37 | pTSAR | | This study | This study |
| *S. flexneri* 2457T ΔipaC | pBAD33 | pMEIC | | This study | This study |
| *S. flexneri* 2457T ΔipaC | pBAD33-ipaC-FLAG37 | pMEIC | | This study | This study |
| *S. flexneri* 2457T ΔipaC | pBAD33-ipaC-FLAG37 | pMEIC-ipaC | | This study | This study |
| *S. flexneri* 2457T ΔipaC | pBAD33-ipaC-FLAG37 | pMEIC-ipaC-A354P | | This study | This study |
| *S. flexneri* 2457T ΔipaC | pBAD33-ipaC-FLAG37 | pMEIC-ipaC-Q308P | | This study | This study |
| *S. flexneri* 2457T ΔipaC | pBAD33-ipaC-FLAG37 | pMEIC-ipaC-ΔC-terminus | | This study | This study |
| *S. flexneri* 2457T ΔipaC | pBAD33-ipaC-FLAG37 | pMEIC-ipaC-Δcoiled-coil | | This study | This study |
| *S. flexneri* 2457T ΔipaC | pBAD33 | pMEIC | pNG162-Afa-1 | This study | This study |
| *S. flexneri* 2457T ΔipaC | pBAD33-ipaC-FLAG37 | pMEIC | pNG162-Afa-1 | This study | This study |
| *S. flexneri* 2457T ΔipaC | pBAD33-ipaC-FLAG37 | pMEIC-ipaC | pNG162-Afa-1 | This study | This study |
| *S. flexneri* 2457T ΔipaC | pBAD33-ipaC-FLAG37 | pMEIC-ipaC-A354P | pNG162-Afa-1 | This study | This study |
| *S. flexneri* 2457T ΔipaC | pBAD33-ipaC-FLAG37 | pMEIC-ipaC-Q308P | pNG162-Afa-1 | This study | This study |
| *S. flexneri* 2457T ΔipaC | pBAD33-ipaC-FLAG37 | pMEIC-ipaC-ΔC-terminus | pNG162-Afa-1 | This study | This study |
| *S. flexneri* 2457T ΔipaC | pBAD33-ipaC-FLAG37 | pMEIC-ipaC-Δcoiled-coil | pNG162-Afa-1 | This study | This study |
| *S. flexneri* 2457T |  |  |  | Laboratory stock | [41] |
| *S. flexneri* 2457T ΔicsB | | | | Laboratory stock | [21] |
| *S. flexneri* 2457T ΔipgB1 | | | | Laboratory stock | [31] |
| *S. flexneri* 2457T | pBAD33 | pMEIC |  | This study | This study |
| *S. flexneri* 2457T ΔicsB | pBAD33 | pMEIC | | This study | This study |
| *S. flexneri* 2457T ΔipgB1 | pBAD33 | pMEIC | | This study | This study |
| *E. coli* DH10B |  |  |  | Catalog no. 18290015. | Thermo Fisher |
| *E. coli* DH10B | pBAD33-ipaC-FLAG37 |  |  | This study | This study |
| *E. coli* DH10B | pMEIC |  |  | This study | This study |
| *E. coli* DH10B | pMEIC-ipaC |  |  | This study | This study |
| *E. coli* DH10B | pMEIC-ipaC-A354P |  |  | This study | This study |
| *E. coli* DH10B | pMEIC-ipaC-Q308P |  |  | This study | This study |
| *E. coli* DH10B | pMEIC-ipaC-ΔC-terminus | | | This study | This study |
| *E. coli* DH10B | pMEIC-ipaC-Δcoiled-coil | | | This study | This study |

### Cell Culture

HeLa, Caco-2, Mouse Embryotic Fibroblast (MEF) cells, HeLa cells that also stably express plasma membrane-targeted yellow fluorescent protein (pmbYFP)^21^ (gift from Hervé Agaisse), or HT-29 cells that stably expressing pmbYFP^21^ (gift from Hervé Agaisse), were grown in Dulbecco’s modified Eagle’s Medium (DMEM) supplemented with 10% fetal bovine serum (FBS) and maintained at 37°C in 5% CO_2_. All cell lines are periodically tested for mycoplasma.

### Fractionation of Infected Cells

HeLa or Caco-2 cells were seeded at 4 x 10^5^ cells per well in a six-well plate. Bacteria were cultured at 37°C at 250 rpm overnight, then back diluted with 1.2% arabinose. Two wells of cells per condition were infected with bacteria at an MOI of 200. To enhance the efficiency of infection, bacteria expressed the *E. coli* adhesion Afa-1^9, 17, 38^. Bacteria were centrifuged onto cells at 800 x *g* for 10 minutes before incubation at 37°C in 5% CO_2_. 1.2% arabinose was or was not included in the media throughout infection to control i*paC* expression from the pBAD promoter during infections with *S. flexneri ΔipaC*::pBAD33- *ipaC* + pTSAR. To only test IpaC variants at spread, arabinose was only included in the bacterial culture media prior to infections with *Sf ΔipaC*::pBAD33-*ipaC-FLAG37* + pMEIC- *ipaC* variant strains. After 45 minutes, cells were washed and remaining extracellular bacteria were killed by addition of 25 μg/mL gentamicin. Cells were fractionated at the indicated timepoints using detergents, as done previously^8–10, 17–20^. In brief, cells were washed three times with ice cold 150 mM Tris, pH 7.4 and scraped into 150 mM Tris, pH 7.4 containing protease inhibitors (protease inhibitor cocktail, complete mini-EDTA free; Roche). Scraped cells were collected and pelleted at 3,000 x *g* for 3.5 min at room temp. The pelleted cells were washed once, resuspended in 150 mM Tris, pH 7.4 containing protease inhibitors and 0.2% saponin, and incubated on ice for 20 min. The suspension was centrifuged at 16,900 x *g* for 30 min at 4°C. The supernatant, which contains the cytosol fraction, was transferred to a fresh tube. The pellet was resuspended in 150 mM Tris, pH 7.4 containing protease inhibitors and 0.5% Triton X-100, incubated on ice for 30 min, and centrifuged at 16,900 x *g* for 15 min at 4°C. The supernatant from this spin contained the membrane fraction, and the pellet consisted of the detergent-insoluble fraction, which included intact bacteria. The abundance of IpaC in the membrane and other fractions was determined by western blot.

### Western Blot

Samples were run on SDS-PAGE gels and transferred to nitrocellulose using a TurboBlot semi dry transfer apparatus (BioRad). The following antibodies were used for western blots: rabbit anti-IpaC^39^ (diluted 1:10,000, incubated 2 hours at room temperature, gift from Wendy Picking), mouse anti-IpaB^40^ (1:15,000, 2 hours at room temperature, gift from Robert Kaminski WRAIR), rabbit anti-GroEL (1:1,000,000, 2 hours at room temperature, catalog no. G6352; Sigma), rabbit anti-caveolin-1 (1:1,000, 2 hours at room temperature, catalog no. C4490; Sigma), mouse anti-E-cadherin (1:500, overnight at 4°C, catalog no. 14-3249-82; Invitrogen), goat anti-rabbit conjugated with horseradish peroxidase (HRP) (1:5,000, 2 hours at room temperature, Jackson ImmunoResearch, catalog no. 115-035- 144), goat anti-mouse conjugated with HRP (1:5,000, 2 hours at room temperature, catalog no. 111-035-146; Jackson ImmunoResearch), goat anti-rat conjugated with HRP (1:5,000, 2 hours at room temperature, catalog no. 115-035-143; Jackson ImmunoResearch). Western blots were developed with Thermo Scientific SuperSignal West Pico and Femto PLUS Chemiluminescent Substrate, and a G:Box (Syngene) was used to acquire the signal. Band intensity was measured using ImageJ (NIH) with the background subtracted.

### Infection and Fluorescence Staining

HeLa pmbYFP or HT-29 pmbYFP cells were seeded at 4 x 10^5^ or 1 x 10^7^ cells, respectively, per well on coverslips in a six-well plate. Bacteria were cultured at 37°C at 250 rpm overnight, then back diluted with 1.2% arabinose. Cells were infected with bacteria at an MOI of 200. Bacteria were centrifuged onto cells at 800 x *g* for 10 minutes before incubation at 37°C in 5% CO_2_. 1.2% arabinose was or was not included in the media throughout infection to control i*paC* expression from the pBAD promoter during infections with *Sf ΔipaC*::pBAD33-*ipaC* + pTSAR. To only test IpaC variants at spread, arabinose was only included in the bacterial culture media prior to infections with *Sf ΔipaC*::pBAD33-*ipaC-FLAG37* + pMEIC-*ipaC* variant strains. After 45 minutes, cells were washed and remaining extracellular bacteria were killed by addition of 25 μg/mL gentamicin. At the indicated timepoints, cells were washed and fixed with 4% paraformaldehyde. If staining for Rac1, cells were permeabilized with 1% triton X-100 for 15 minutes, then washed and stained with anti-Rac1 (1:50, overnight at 4°C, catalog no. 610650; BD Biosciences) before being washed and incubated with goat anti-mouse Alexa-Fluor 750 (1:100, 2 hours at 25°C, catalog no. A21037; Invitrogen) for 2 hours at 25°C. All fixed infected cells were washed, stained with Hoechst, and the coverslips were mounted on slides with ProLong Diamond.

### Microscopy

Fluorescence images were acquired on a Nikon Eclipse Ti-2 inverted light microscope equipped with an Orca Fusion BT cMOS camera (Hammamatsu) and Semrock Brightline filters. Images were collected at 100X magnification randomly across the coverslip using an automated imaging pipeline created in NIS elements software (Nikon). Images were deconvolved using a Richardson-Lucy algorithm with 20 iterations. Microscopic images were pseudo-colored and assembled using FIJI (NIH). Z-stacks with twenty-seven slices of 0.20 μm step size were collected per position on a coverslip. To display representative images, slices within a stack were collapsed into a single image based upon the maximum intensity of pixels in each slice using NIS elements software. NIS elements software was used to identify GFP positive and/or mCherry positive bacteria. Bacteria within DMVs were classified by their complete enclosure within a YFP membrane and with no membrane tether as defined previously^21, 30^.

### Plaque Assays

MEFs were seeded in six-well plates at 6 x 10^6^ cells per well. Bacteria were cultured at 37°C at 250 rpm overnight, then back diluted with 1.2% arabinose. Cells were infected with bacteria at an MOI of 0.02 with or without 1.2% arabinose in the cell culture media. Bacteria were centrifuged onto cells at 800 x *g* for 10 minutes and incubated at 37°C in 5% CO_2_ for 1 hour. Media was replaced with DMEM containing 10% FBS, 25 μg/mL gentamicin, 1.2% arabinose where indicated, and 0.5% agarose, and the cells were incubated for 48 hours before an overlay of 0.7% agarose in DMEM with 10% FBS, 25 μg/mL gentamicin, and 0.1% neutral red was added. After 4-6 hours of incubation, plaques were imaged with an ImmunoSpot S6 Universal Visible/Fluorescent Analyzer (Cellular Technology Limited). Plaques were analyzed with a pipeline generated in ImageJ in which images were thresholded to remove background, pixel intensity was converted to a binary, and the images were segmented into objects; objects matching plaques on unmodified images were selected, and their area was quantified, as done previously^10^.

### Statistical Analysis

Statistical differences between means were determined with GraphPad Prism (version 10). Differences between the means of three or more groups were tested by either two-way ANOVA with Fisher’s or Sidak’s *post hoc* test or one-way ANOVA with Dunnett’s *post hoc* test. Differences between the means of two groups were tested by Student’s t test.

## ACKNOWLEDGEMENTS

We would like to thank Robert Kaminski and Hervé Agaisse for reagents. We would like to thank Cristina Penaranda, Lilian Radoshevich, Marijke Keestra-Gounder and the Keestra-Gounder lab for helpful discussions. This work was funded by K22 AI137296, the GI and Liver Innate Immune Program at the University of Colorado, and the University of Colorado start-up funds to BCR. JER was funded by T32 AI052066.

## Notes

### Competing Interest Statement

The authors have declared no competing interest.

