## Supplemental Material for "A translocation-competent pore is required for *Shigella flexneri* to escape from the double membrane vacuole during intercellular spread"

Running Head: Translocation-competent pores enable *S.flexneri* spread.

### SUPPLEMENTARY MATERIALS AND METHODS

#### Secretion Assays

Bacteria were cultured at 37°C with shaking at 250 rpm overnight, then back diluted, induced when indicated with 1.2% arabinose, and normalized by measuring optical density at 600 nm ($OD_{600}$). Bacteria were resuspended in phosphate-buffered saline (PBS) containing 1.2% arabinose, where appropriate, and 10 µM Congo red where indicated, and incubated for 2 hours in a 37°C water bath. Bacteria were pelleted at 15,000 x *g* and the supernatant collected. Western blots were used to detect IpaC and IpaC-FLAG37 in the pellets and supernatants.

#### Pore formation by erythrocyte lysis assay

Pore formation in sheep erythrocyte membranes was monitored by assessing the efficiency of erythrocyte lysis as done previously[1-4]. Briefly, defibrinated sheep erythrocytes (HemoStat) were pelleted at 2,000 x *g* and resuspended in 100 µL of PBS. Erythrocytes were infected at an MOI of 25 in 100 µl of PBS supplemented with 1.2% arabinose. Bacteria were centrifuged onto the erythrocytes at 2,000 x *g* for 10 min at 25°C and were co-cultured with the erythrocytes for 50 minutes at 37°C. The bacterial and erythrocyte cocultures were mixed by pipetting and then centrifuged again at 2,000 x *g* for 10 min at 25°C. As a positive control for lysis, an aliquot of uninfected erythrocytes was treated with 0.02% SDS. The supernatants were collected, and abundance of hemoglobin released was measured by absorbance at 570 nm using an Epoch II plate reader (BioTech) or Wallac 1420 Victor2 microplate reader (Perkin Elmer).

**Translocation and docking**

The measurement of docking and T3SS activation was performed as previously described[1, 2, 4, 5]. Briefly, MEFs were seeded at 3 x $10^5$ cells per well on coverslips in a six-well plate. Bacteria that constitutively produce mCherry under the rpsM promoter and harbor the TSAR reporter plasmid were grown to exponential phase and added to cells at an MOI of 200. The TSAR reporter, which expresses green fluorescent protein (GFP) when the bacterial effector OspD is secreted through the T3SS[6], is an indicator of active T3SS secretion. Bacteria were then centrifuged onto cells at 800 x *g* for 10 minutes at 25°C. The co-culture was incubated at 37°C for an additional 50 minutes. The infected cells were washed with HBSS and fixed with 3.7% paraformaldehyde. DNA was stained with Hoechst. Coverslips were mounted onto glass slides with ProLong Diamond (Invitrogen). Bacteria were examined by fluorescence microscopy using a Nikon Eclipse TE-300 with appropriate filters. Bacterial docking was quantified by determining the number of mCherry-producing bacteria that remained associated with cells. Bacteria with active T3SS were determined by counting the number of cell-associated bacteria expressing GFP. For each condition of each independent experiment, 20 to 250 eukaryotic cells and 20 to 210 bacteria were analyzed.

### SUPPLEMENTARY FIGURE LEGENDS

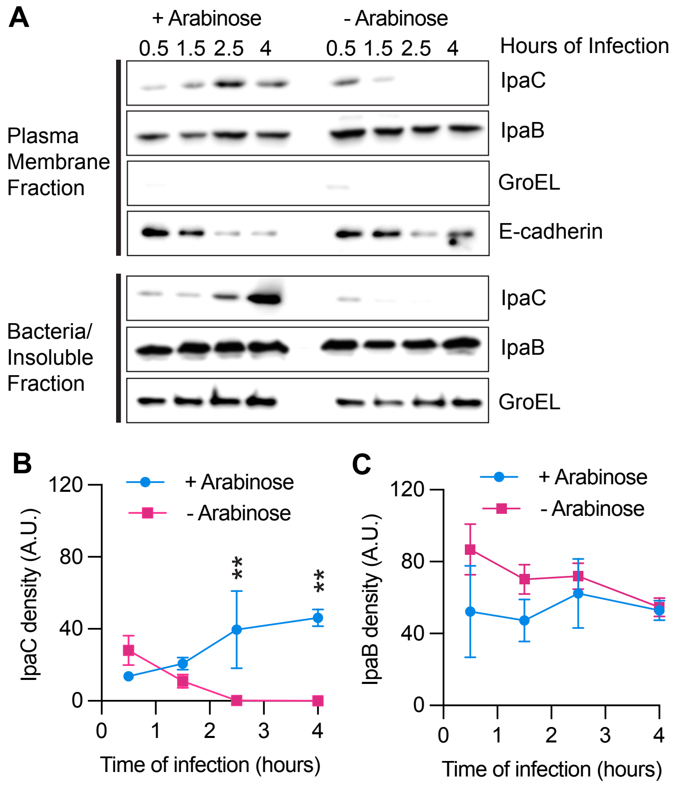


**Supplementary Figure 1: IpaC and IpaB are present in the host membrane during infection of Caco-2 cells.**

**A)** Representative western blots showing abundance of IpaC, IpaB, GroEL (bacterial cytoplasmic protein), and E-cadherin (host membrane protein) in the membrane fraction (top panels) and of IpaC, IpaB, and GroEL in the bacteria/insoluble fraction (bottom panels) from infected Caco-2 cells. **B & C)** Quantification of the amount of IpaC (**B)** or IpaB (**C)** in the membrane fraction with (light blue) or without (magenta) arabinose included in the media during infection to induce IpaC production. Three independent experiments were performed for each infection condition at each timepoint. Data are mean ± SEM, some error bars are smaller than the symbol and are not visible. Mixed effects analysis with Fisher's multiple comparisons test, **$p<0.01$.

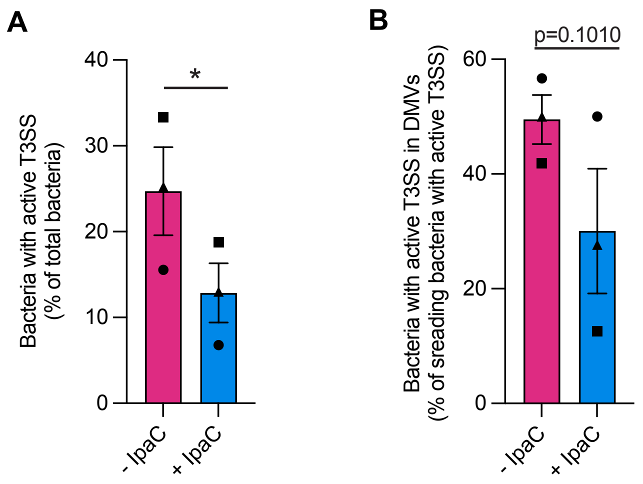


**Supplementary Figure 2**: **Bacteria not producing IpaC during spread show more T3SS activation.**

**A)** Percentage of bacteria with active T3SS at 4 hours of infection from images represented in Fig. 2A. **B)** Percentage of bacteria that are spreading and in DMVs with active T3SS. **A & B)** Bacteria producing (+ IpaC, aqua) or not producing (- IpaC, magenta) IpaC during infection. Data are mean ± SEM of three independent experiments, each experiment matched by symbol. *$p<0.05$, by paired t-test; ns (not significant).

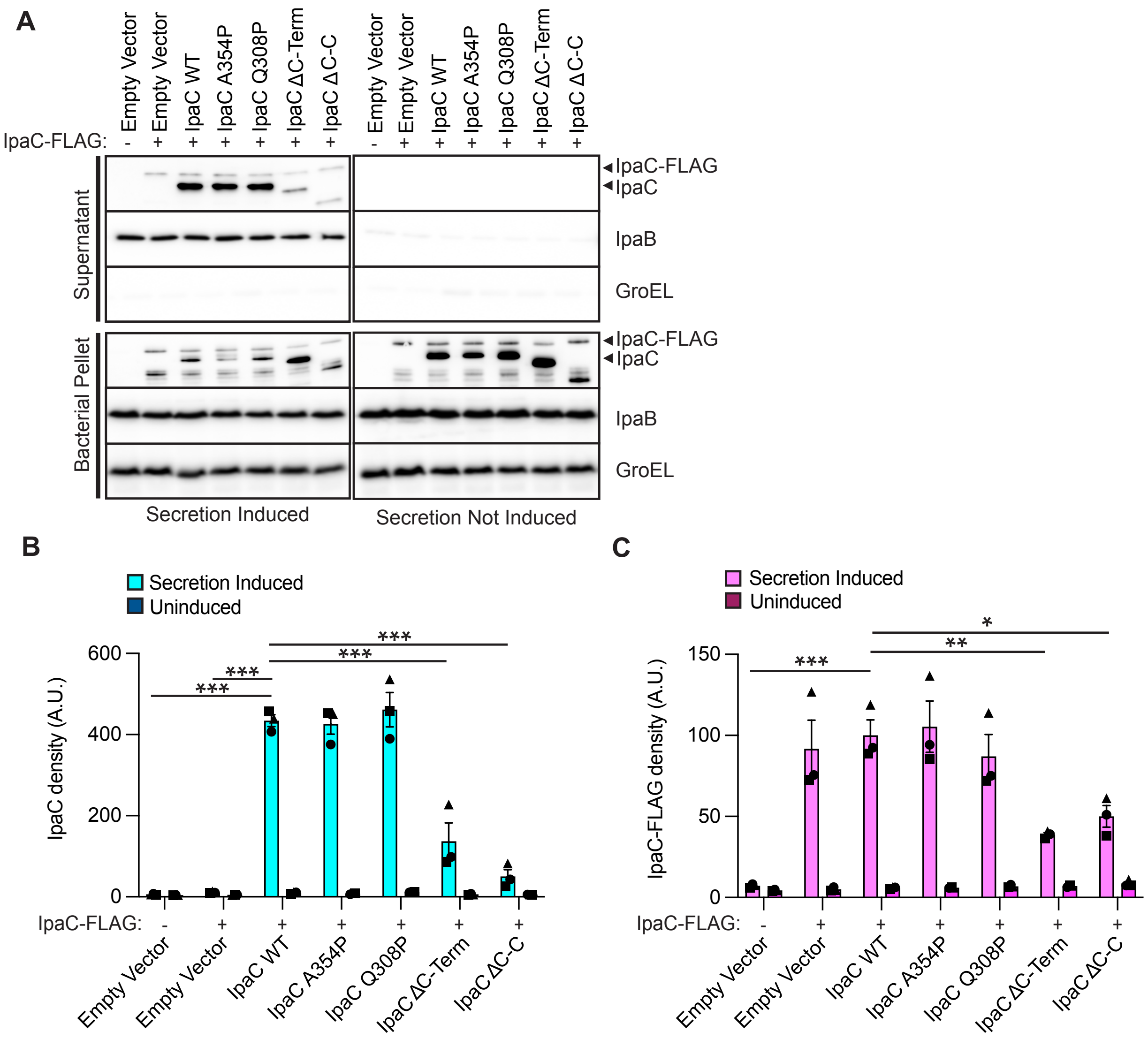


**Supplementary Figure 3**: ***S. flexneri* producing IpaC variants and IpaC-FLAG can secrete both when induced.**

**A)** Representative western blots showing abundance of IpaC-FLAG (induced), IpaC variants, IpaB, and GroEL in the supernatant (top panels) and the bacterial pellet (bottom panels) following induction or not of secretion by Congo red. C-Term (C-terminus), C-C (Coiled-coil). **B & C)** Quantification of the amount of IpaC variants **(B)** or IpaC-FLAG **(C)** in the supernatant when secretion is induced or uninduced. Data are mean ± SEM of three independent experiment, each experiment matched by symbol. Two-way ANOVA with Dunnett's multiple comparisons test; ns (not significant), *p<0.05, **p<0.01, ***p<0.001.

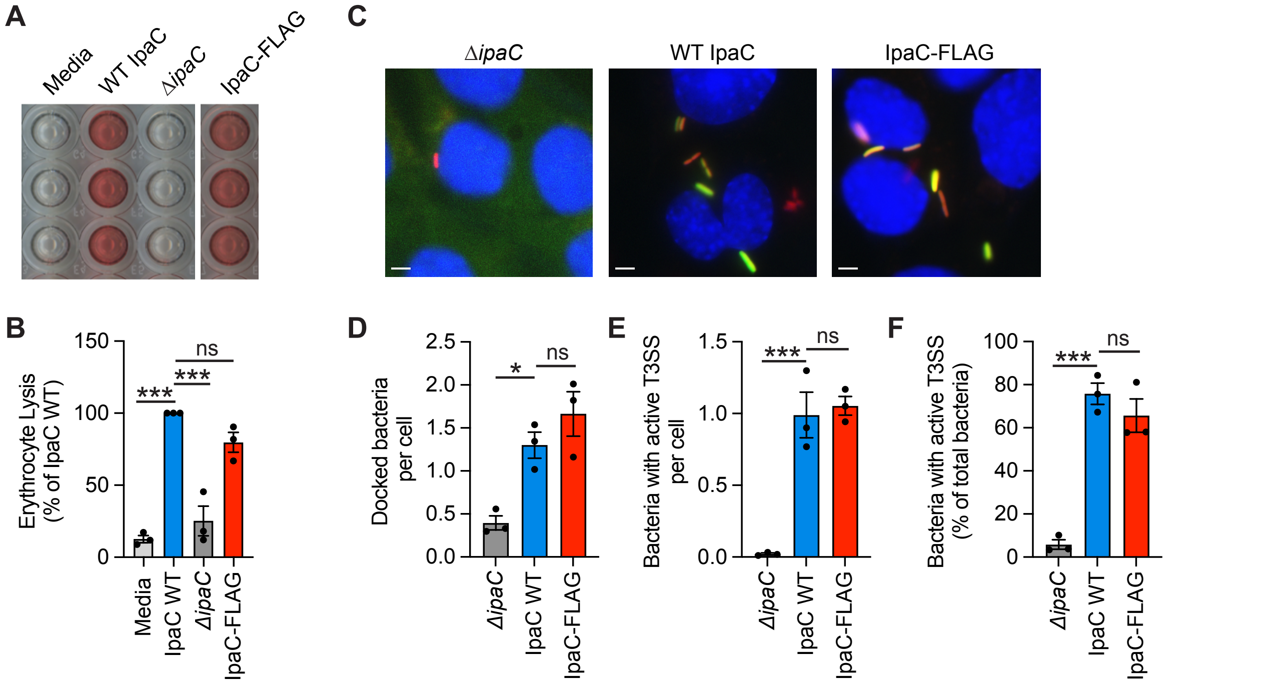


**Supplementary Figure 4**: ***S. flexneri*** **producing IpaC-FLAG can form pores and translocate effectors.**

**A)** Representative image of hemoglobin released from erythrocytes. **B)** Quantification of hemoglobin released in experiments represented in A. Data points are individual experiments; data are the mean ± SEM of three experiments per strain. N.S., not significant; ***, p<0.001. One-way ANOVA with Dunnett's multiple comparisons test. **C)** Representative images of infection of MEFs by bacteria producing no IpaC, IpaC, or IpaC-FLAG after 1 hour of infection. Red, bacteria; green, bacteria with active T3SS; blue, DNA; (scale bar 5 µm. Number of docked bacteria per cell **(D)**, number of bacteria with active T3SS secretion per cell **(E)**, and percentage of docked bacteria with activate T3SS secretion **(F)** in experiments represented in panel **C. D-F)** Data points are individual experiments; data are the mean ± SEM of three experiments per strain. N.S., not significant; *, p<0.05; ***, p<0.001. One-way ANOVA with Dunnett's multiple comparisons test.

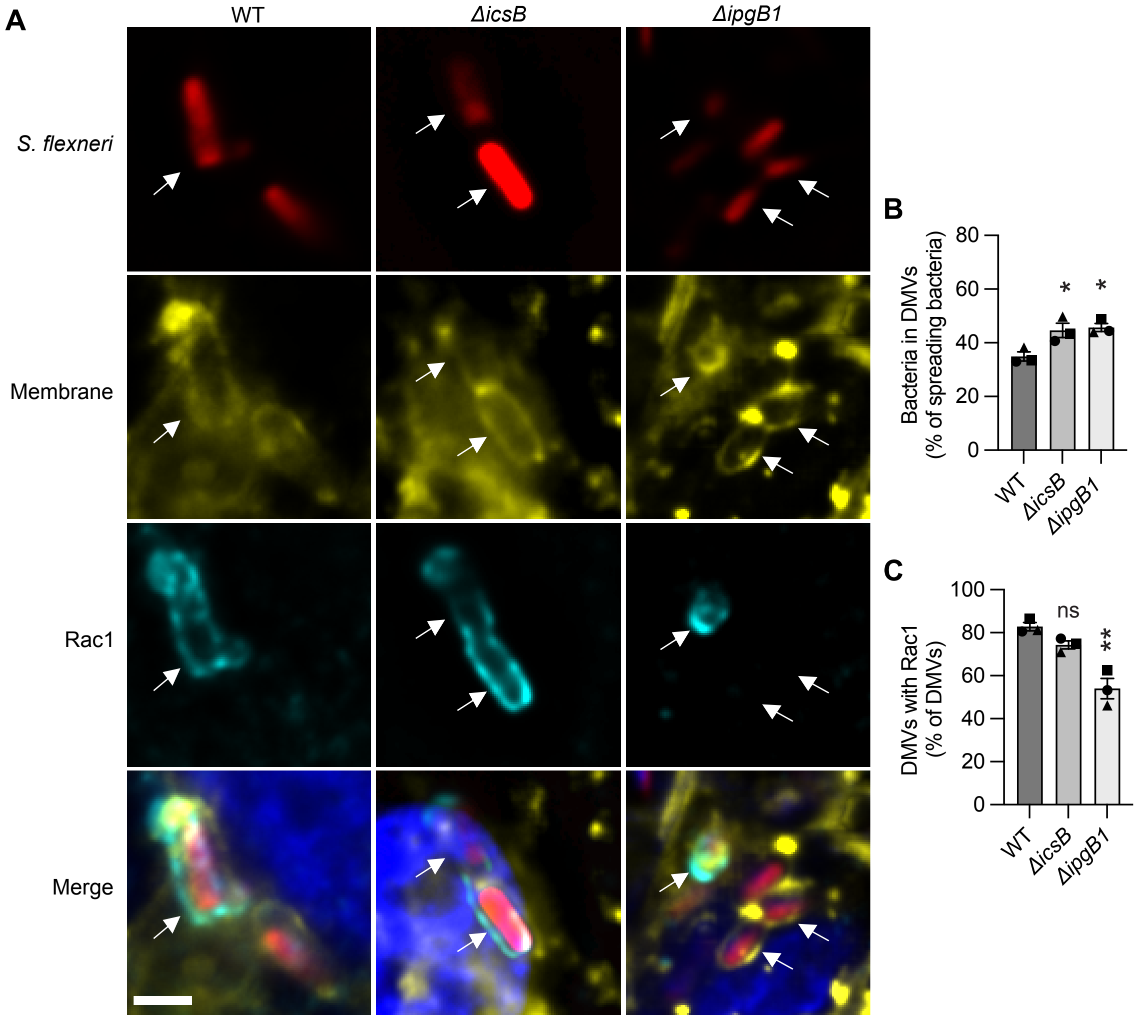


**Supplementary Figure 5: Rac1 colocalization with DMVs is associated with IpgB1 function.**

**A)** Representative immunofluorescence images of HT-29 pmbYFP cells infected with bacteria producing indicated *S. flexneri* knockout at 5 hours of infection. Bacteria in a DMV (white arrow). Red, bacteria; yellow, HT-29 cell membranes; cyan, Rac1; blue, DNA;

scale bar, 2 μm. From images represented in **A,** percent of bacteria that are spreading and in DMVs **(B)**, and percent of bacteria within DMVs that colocalize with Rac1 (**C)**. **B-C)** Data are mean ± SEM of three independent experiments, each experiment matched by symbol. ns (not significant), *p<0.05, **p<0.01 by one-way ANOVA with Dunnett's multiple comparisons test.

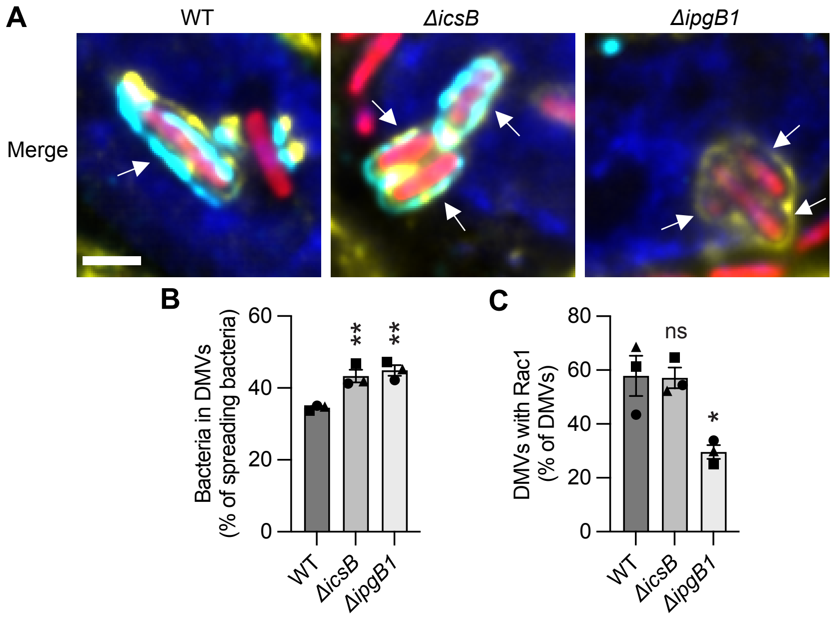


**Supplementary Figure 6: Rac1 colocalization with DMVs is associated with IpgB1 function in HeLa cells.**

**A)** Representative immunofluorescence images of HeLa pmbYFP cells infected with bacteria producing indicated *S. flexneri* knockout at 4 hours of infection. Bacteria within a DMV (white arrow). Red, bacteria; yellow, HeLa cell membranes; cyan, Rac1; blue, DNA; scale bar, 2 μm. From images represented in **A,** percent of bacteria that are spreading and in DMVs **B)**, and percent of bacteria within DMVs that colocalize with Rac1 **C)**. **B-C)** Data are mean ± SEM of three independent experiments, each experiment matched by symbol. ns (not significant), *p<0.05, **p<0.01 by one-way ANOVA with Dunnett's multiple comparisons test.

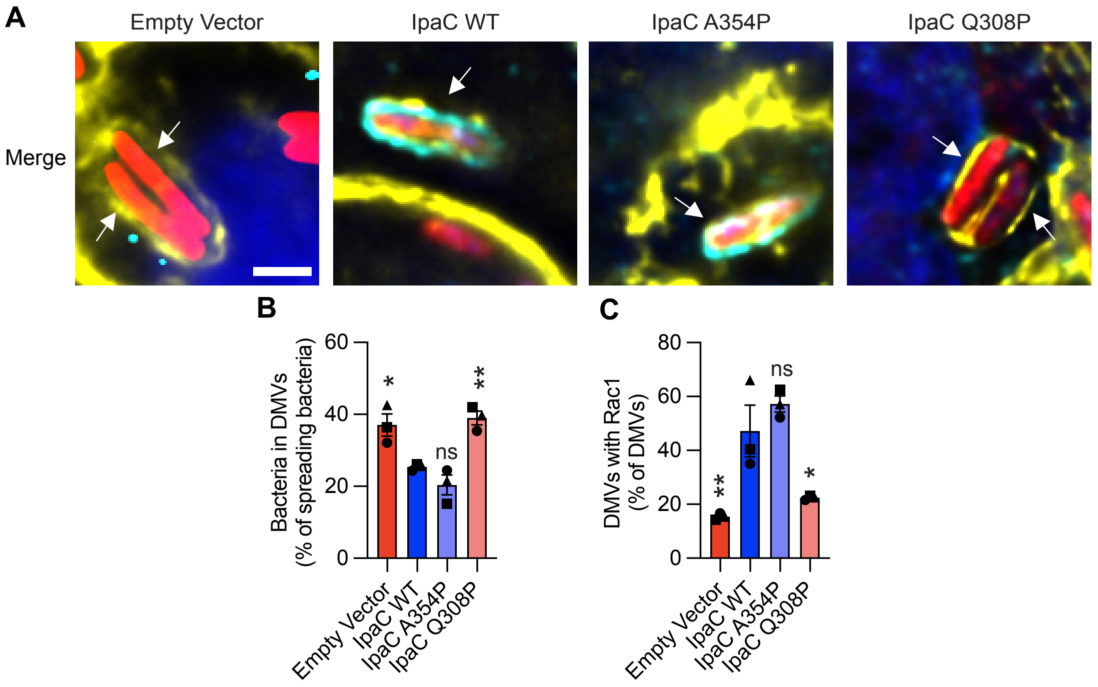


**Supplementary Figure 7**: **Translocation-competent pores are required for *S. flexneri* to recruit Rac1 to DMVs for efficient escape in HeLa cells.**

**A)** Representative immunofluorescence images of HeLa pmbYFP cells infected with bacteria producing indicated IpaC variants at 4 hours of infection. All strains included IpaC-FLAG and were induced only before infection to enable invasion. Bacteria within a DMV (white arrow). Red, bacteria; yellow, HeLa cell membranes; cyan, Rac1; blue, DNA; scale bar, 2 μm. From images represented in **A, B)** percent of bacteria within DMVs that colocalize with Rac1 and **C)** percent of bacteria that are spreading and in DMVs. **B & C)** Data are mean ± SEM of three independent experiments, each experiment matched by symbol. ns (not significant), *p<0.05, **p<0.01 by one-way ANOVA with Dunnett's multiple comparisons test.

| | Total bacteria | | Spreading bacteria (% of total bacteria) | | Bacteria in DMVs (% of total bacteria) | | Total bacteria in DMVs (% of spreading bacteria) | | Bacteria with active T3SS (% of total bacteria) | | Spreading bacteria with active T3SS (% of bacteria with active T3SS) | | Bacteria with active T3SS in DMVs (% of bacteria with active T3SS) | | Bacteria with active T3SS in DMVs (% of spreading bacteria with active T3SS) | |
|---|---|---|---|---|---|---|---|---|---|---|---|---|---|---|---|---|
| | **- IpaC** | **+ IpaC** | **- IpaC** | **+ IpaC** | **- IpaC** | **+ IpaC** | **- IpaC** | **+ IpaC** | **- IpaC** | **+ IpaC** | **- IpaC** | **+ IpaC** | **- IpaC** | **+ IpaC** | **- IpaC** | **+ IpaC** |
| Experiment 1 | 765 | 942 | 83 (10.9%) | 70 (7.4%) | 32 (4.2%) | 22 (2.3%) | 32 (38.6%) | 22 (31.4%) | 119 (15.6%) | 64 (6.8%) | 30 (36.1%) | 28 (40.0%) | 17 (14.3%) | 14 (21.9%) | 17 (56.7%) | 14 (50.0%) |
| Experiment 2 | 2755 | 2330 | 391 (14.2%) | 386 (16.6%) | 129 (4.7%) | 75 (3.2%) | 129 (33.0%) | 75 (19.4%) | 918 (33.3%) | 437 (18.8%) | 172 (44.0%) | 223 (57.8%) | 72 (7.8%) | 28 (6.4%) | 72 (41.9%) | 28 (12.6%) |
| Experiment 3 | 1192 | 1260 | 186 (15.6%) | 180 (14.3%) | 77 (6.5%) | 50 (4.0%) | 77 (41.4%) | 50 (24.8%) | 301 (25.3%) | 164 (13.0%) | 52 (28.0%) | 47 (26.1%) | 26 (8.6%) | 13 (7.9%) | 26 (50.0%) | 13 (27.7%) |

**Supplementary Table 1: Quantifications of bacteria from Figure 2.**

Quantification of images acquired across three independent experiments at 4 hours of infection in Figure 2 for bacteria not producing (- IpaC, pink) or producing (+ IpaC, blue) IpaC during infection. Total bacteria: all mCherry+ bacteria. Spreading bacteria: all mCherry+ bacteria observed in protrusions, vacuole-like protrusions, or DMVs. Bacteria in DMVs: all mCherry+ bacteria observed in DMVs. Bacteria with active T3SS: mCherry+ bacteria that are also GFP+. Spreading bacteria with active T3SS: Bacteria with active T3SS observed in protrusions, vacuole-like protrusions, or DMVs. Bacteria with active T3SS in DMVs: Bacteria with active T3SS observed in DMVs.

| | Total bacteria | | | Spreading bacteria (% of total bacteria) | | | Bacteria in DMVs (% of total bacteria) | | | Bacteria in DMVs (% of spreading bacteria) | | | DMVs with Rac1 (% of DMVs) | | |
|---|---|---|---|---|---|---|---|---|---|---|---|---|---|---|---|
| | WT | *ΔicsB* | *ΔipgB1* | WT | *ΔicsB* | *ΔipgB1* | WT | *ΔicsB* | *ΔipgB1* | WT | *ΔicsB* | *ΔipgB1* | WT | *ΔicsB* | *ΔipgB1* |
| Experiment 1 | 495 | 1172 | 527 | 109 (22.0%) | 248 (21.2%) | 169 (32.1%) | 36 (7.3%) | 101 (8.6%) | 75 (14.2%) | 36 (33.0%) | 101 (40.7%) | 75 (44.4%) | 29 (80.6%) | 78 (77.2%) | 40 (53.3%) |
| Experiment 2 | 1018 | 1364 | 1887 | 245 (24.1%) | 311 (22.8%) | 367 (19.4%) | 82 (8.1%) | 135 (9.9%) | 179 (9.5%) | 82 (33.5%) | 135 (43.4%) | 179 (48.8%) | 71 (86.6%) | 101 (74.8%) | 112 (62.6%) |
| Experiment 3 | 1680 | 1523 | 829 | 392 (23.3%) | 187 (12.3%) | 182 (22.0%) | 150 (8.9%) | 93 (6.1%) | 80 (9.7%) | 150 (38.2%) | 93 (49.7%) | 80 (44.0%) | 122 (81.3%) | 66 (71.0%) | 37 (46.3%) |

**Supplementary Table 2: Quantifications of bacteria from infected HT-29 cells in Supplementary Figure 5.**

Quantification of images acquired across three independent experiments at 5 hours of infection in Supplementary Figure 5 for indicated *S. flexneri* knockouts. Total bacteria: all mCherry+ bacteria. Spreading bacteria: all mCherry+ bacteria observed in protrusions, vacuole-like protrusions, or DMVs. Bacteria in DMVs: all mCherry+ bacteria observed in DMVs. Bacteria in DMVs with Rac1: all mCherry+ bacteria observed in DMVs that also colocalize with Rac1.

| | Total bacteria | | | Spreading bacteria (% of total bacteria) | | | Bacteria in DMVs (% of total bacteria) | | | Bacteria in DMVs (% of spreading bacteria) | | | DMVs with Rac1 (% of DMVs) | | |
|---|---|---|---|---|---|---|---|---|---|---|---|---|---|---|---|
| | WT | Δ*icsB* | Δ*ipgB1* | WT | Δ*icsB* | Δ*ipgB1* | WT | Δ*icsB* | Δ*ipgB1* | WT | Δ*icsB* | Δ*ipgB1* | WT | Δ*icsB* | Δ*ipgB1* |
| Experiment 1 | 352 | 860 | 492 | 131 (37.2%) | 165 (19.2%) | 154 (31.3%) | 46 (13.1%) | 68 (7.9%) | 65 (13.2%) | 46 (35.1%) | 68 (41.2%) | 65 (42.2%) | 20 (43.5%) | 37 (54.4%) | 22 (33.8%) |
| Experiment 2 | 1013 | 855 | 713 | 169 (16.7%) | 109 (12.7%) | 144 (20.2%) | 57 (5.6%) | 51 (6.0%) | 68 (9.5%) | 57 (33.7%) | 51 (46.8%) | 68 (47.2%) | 35 (61.4%) | 33 (64.7%) | 17 (25.0%) |
| Experiment 3 | 1904 | 1540 | 1623 | 247 (13.0%) | 210 (13.6%) | 325 (20.0%) | 86 (4.5%) | 88 (5.7%) | 147 (9.1%) | 86 (34.8%) | 88 (41.9%) | 147 (45.2%) | 59 (68.6%) | 46 (52.3%) | 44 (29.9%) |

**Supplementary Table 3: Quantifications of bacteria from infected HeLa cells in Supplementary Figure 6.**

Quantification of images acquired across three independent experiments at 4 hours of infection in Supplementary Figure 6 for indicated *S. flexneri* knockouts. Total bacteria: all mCherry+ bacteria. Spreading bacteria: all mCherry+ bacteria observed in protrusions, vacuole-like protrusions, or DMVs. Bacteria in DMVs: all mCherry+ bacteria observed in DMVs. Bacteria in DMVs with Rac1: all mCherry+ bacteria observed in DMVs that also colocalize with Rac1.

| | Total bacteria | | | | Spreading bacteria (% of total bacteria) | | | | Bacteria in DMVs (% of total bacteria) | | | | Bacteria in DMVs (% of spreading bacteria) | | | | DMVs with Rac1 (% of DMVs) | | | |
|---|---|---|---|---|---|---|---|---|---|---|---|---|---|---|---|---|---|---|---|---|
| | Empty Vector | IpaC WT | IpaC A354P | IpaC Q308P | Empty Vector | IpaC WT | IpaC A354P | IpaC Q308P | Empty Vector | IpaC WT | IpaC A354P | IpaC Q308P | Empty Vector | IpaC WT | IpaC A354P | IpaC Q308P | Empty Vector | IpaC WT | IpaC A354P | IpaC Q308P |
| Experiment 1 | 202 | 522 | 716 | 124 | 94 (46.5%) | 90 (17.2%) | 183 (25.6%) | 49 (39.5%) | 57 (28.2%) | 31 (5.9%) | 64 (8.9%) | 25 (20.2%) | 57 (60.6%) | 31 (34.4%) | 64 (35.0%) | 25 (51.0%) | 2 (3.5%) | 15 (48.4%) | 47 (73.4%) | 0 (0.0%) |
| Experiment 2 | 237 | 943 | 340 | 176 | 87 (36.7%) | 168 (17.8%) | 93 (27.4%) | 60 (34.1%) | 40 (16.9%) | 48 (5.1%) | 25 (7.4%) | 26 (14.8%) | 40 (46.0%) | 48 (28.6%) | 25 (26.9%) | 26 (43.3%) | 7 (17.5%) | 37 (77.1%) | 21 (84.0%) | 8 (30.8%) |
| Experiment 3 | 421 | 475 | 631 | 332 | 114 (27.1%) | 101 (21.3%) | 162 (25.7%) | 110 (33.1%) | 56 (13.3%) | 29 (6.1%) | 60 (9.5%) | 59 (17.8%) | 56 (49.1%) | 29 (28.7%) | 60 (37.0%) | 59 (53.6%) | 5 (8.9%) | 23 (79.3%) | 42 (70.0%) | 7 (11.9%) |

**Supplementary Table 4: Quantifications of bacteria from infected HT-29 cells in Figure 5.**

Quantification of images acquired across three independent experiments at 5 hours of infection in Figure 5 for *S. flexneri* producing indicated IpaC variant. Total bacteria: all mCherry+ bacteria. Spreading bacteria: all mCherry+ bacteria observed in protrusions, vacuole-like protrusions, or DMVs. Bacteria in DMVs: all mCherry+ bacteria observed in DMVs. Bacteria in DMVs with Rac1: all mCherry+ bacteria observed in DMVs that also colocalize with Rac1.

| | Total bacteria | | | | Spreading bacteria (% of total bacteria) | | | | Bacteria in DMVs (% of total bacteria) | | | | Bacteria in DMVs (% of spreading bacteria) | | | | DMVs with Rac1 (% of DMVs) | | | |
|---|---|---|---|---|---|---|---|---|---|---|---|---|---|---|---|---|---|---|---|---|
| | Empty Vector | IpaC WT | IpaC A354P | IpaC Q308P | Empty Vector | IpaC WT | IpaC A354P | IpaC Q308P | Empty Vector | IpaC WT | IpaC A354P | IpaC Q308P | Empty Vector | IpaC WT | IpaC A354P | IpaC Q308P | Empty Vector | IpaC WT | IpaC A354P | IpaC Q308P |
| Experiment 1 | 883 | 722 | 1715 | 966 | 56 (6.3%) | 82 (11.4%) | 94 (5.5%) | 144 (14.9%) | 18 (2.0%) | 20 (2.8%) | 23 (1.3%) | 51 (5.3%) | 18 (32.1%) | 20 (24.4%) | 23 (24.5%) | 51 (35.4%) | 3 (16.7%) | 7 (35.0%) | 12 (52.2%) | 11 (21.6%) |
| Experiment 2 | 299 | 446 | 433 | 268 | 77 (25.8%) | 161 (36.1%) | 138 (31.9%) | 124 (46.3%) | 28 (9.4%) | 42 (9.4%) | 32 (7.4%) | 52 (19.4%) | 28 (36.4%) | 42 (26.1%) | 32 (23.2%) | 52 (41.9%) | 4 (14.3%) | 17 (40.5%) | 20 (62.5%) | 12 (23.1%) |
| Experiment 3 | 286 | 1155 | 1297 | 185 | 61 (21.3%) | 242 (21.0%) | 130 (10.0%) | 78 (42.1%) | 26 (9.1%) | 62 (5.4%) | 28 (2.2%) | 31 (16.8%) | 26 (42.6%) | 62 (25.6%) | 28 (21.5%) | 31 (39.7%) | 4 (15.4%) | 41 (66.1%) | 16 (57.1%) | 7 (22.6%) |

**Supplementary Table 5: Quantifications of bacteria from infected HeLa cells in Supplementary Figure 7.**

Quantification of images acquired across three independent experiments at 4 hours of infection in Supplementary Figure 7 for *S. flexneri* producing indicated IpaC variant. Total bacteria: all mCherry+ bacteria. Spreading bacteria: all mCherry+

bacteria observed in protrusions, vacuole-like protrusions, or DMVs. Bacteria in DMVs: all mCherry+ bacteria observed in DMVs. Bacteria in DMVs with Rac1: all mCherry+ bacteria observed in DMVs that also colocalize with Rac1.
